# Single-axon level automatic segmentation and feature extraction from immuhistochemical images of peripheral nerves

**DOI:** 10.1101/2020.06.24.169557

**Authors:** Viktor Tóth, Naveen Jayaprakash, Adam Abbas, Ariba Khan, Stavros Zanos, Theodoros P. Zanos

## Abstract

Quantitative descriptions of the morphology and structure of peripheral nerves is central in the development of bioelectronic devices interfacing the nerves. While histological procedures and microscopy techniques yield high-resolution detailed images of individual axons, automated methods to extract relevant information at the single-axon level are not widely available. We implemented a segmentation algorithm that allows for subsequent feature extraction in immunohistochemistry (IHC) images of peripheral nerves at the single fiber scale. These features include short and long cross-sectional diameters, area, perimeter, thickness of surrounding myelin and polar coordinates of single axons within a nerve or nerve fascicle. We evaluated the performance of our algorithm using manually annotated IHC images of 27 fascicles of the swine cervical vagus; the accuracy of single-axon detection was 82%, and of the classification of fiber myelination was 89%.

## I. Introduction

Interfacing with peripheral nerves is a key component of many bioelectronic medicine devices [1] and brain-machine interfaces (BMI) approaches [2] to treat diseases and conditions. As an example, vagus nerve stimulation is used and tested for drug resistant epilepsy [3], heart failure [4], depression [5], rheumatoid arthritis [6], pulmonary hypertension [7] and tinnitus [8], among others. Despite significant clinical interest, clinical efficacy is often limited and side-effects such as cough, throat-pain, voice alterations and dyspnea regularly occur, since the vagus nerve modulates the function of many organs, via axonal projections of different sizes and distributions. To develop more targeted therapies, we need to describe the functional anatomy of the vagus nerve at the single axon level.

The functional anatomy of the vagus nerve has been studied in various animal models and in humans, using different imaging modalities and tracing techniques [9], [10]. While all these studies provide valuable information with regards to the variability in sizes of the whole nerve and its fascicles, as well as other macroscale characteristics of the nerve, they lack the analysis of features at the level of single fibers, e.g. number of axons, myelination patterns and sizes of fibers.

Advances in immunohistochemistry (IHC) techniques and improvements in the microscopes used to image nerve dissections have enabled acquisition of high quality images that provide not only detailed anatomical features of the different fascicles and nerve fibers, but also visualize discrete cellular components of nerve fibers, including the axon and myelin. However, manual annotation of these features is laborious and cannot easily scale to multiple IHC images of nerves that have dozens of fascicles, containing thousands of myelinated and unmyelinated axons. To obtain quantitative axon-level counts and features, automated methods that will extract this information are needed. To this end, we developed a segmentation and feature extraction algorithm that is applied to IHC images of a peripheral nerve with complex anatomy and translational significance: the cervical vagus nerve of the swine. The algorithm provides information related to counts of myelinated and unmyelinated axons, location, shape and size of individual axons and thickness of the myelin sheath. The algorithm was validated using manually annotated images through which the accuracy of the method was established. The method is not limited to the pig animal model or the vagus nerve, but can be easily extended to other animal models or human IHC images, as well as other peripheral nerves.

## II. Methods

### A. Imaging

Vagus nerve tissue samples were obtained from Yucatan swine postmortem. Standard IHC protocol was followed to stain the sections for neurofilament and myelin basic protein [11]. Briefly, sections were subjected to deparaffinization procedure using xylene and ethanol rinse and then washed with distilled water for 10 min. Citrate antigen retrieval was performed by briefly subjecting the samples to 1 x citrate buffer to unmask the antigens. The sectioned was rinsed, blocked and stained for incubated with Anti-Neurofilament (1:500, Abcam AB8135) and Anti-Myelin basic protein (1:500, Abcam AB7349).

The following day, sections were rinsed and incubated with the secondary antibody (1:500) for two hours at room temperature. Slides were then rinsed thoroughly with TBS buffer for three times and cover glass was mounted on the sides with the fluoromount-G (Thermo Fisher Scientific), while care was taken to avoid air bubbles during the placement of the cover glass. Transparent top coat (Electron Microscopy sciences) was used to secure the cover glass on the sides. Once the polish was dried, the samples were imaged using ZEISS LSM 900, confocal laser scanning microscope.

Across the depth dimension of the scan, the maximum intensity projection was extracted for further analysis. Each of the 27 fascicles from the whole nerve image was cropped out and processed individually. We adjusted the color levels (minimum and maximum levels of 42, 100 and 42, 124 out of 255 for the neurofilament and myelin channels respectively) to increase contrast between the background and the foreground axons. All the 27 fascicles were manually annotated with locational and myelination information for each axon to serve as ground truth in testing our segmentation algorithm.

### B. Segmentation

We developed an algorithm that performs segmentation and feature extraction of IHC images of peripheral nerves. We demonstrate the accuracy of this method on images of the swine vagus nerve. The neurofilament and myelin of the vagus were stained, and they correspond to the green and red channels, respectively (Fig. 1). The segmentation method does not require annotated images to be trained; however, it relies on a few parameters that we manually tuned on images of two fascicles through the visual inspection of resulting segmentations.

**Figure. 1.**
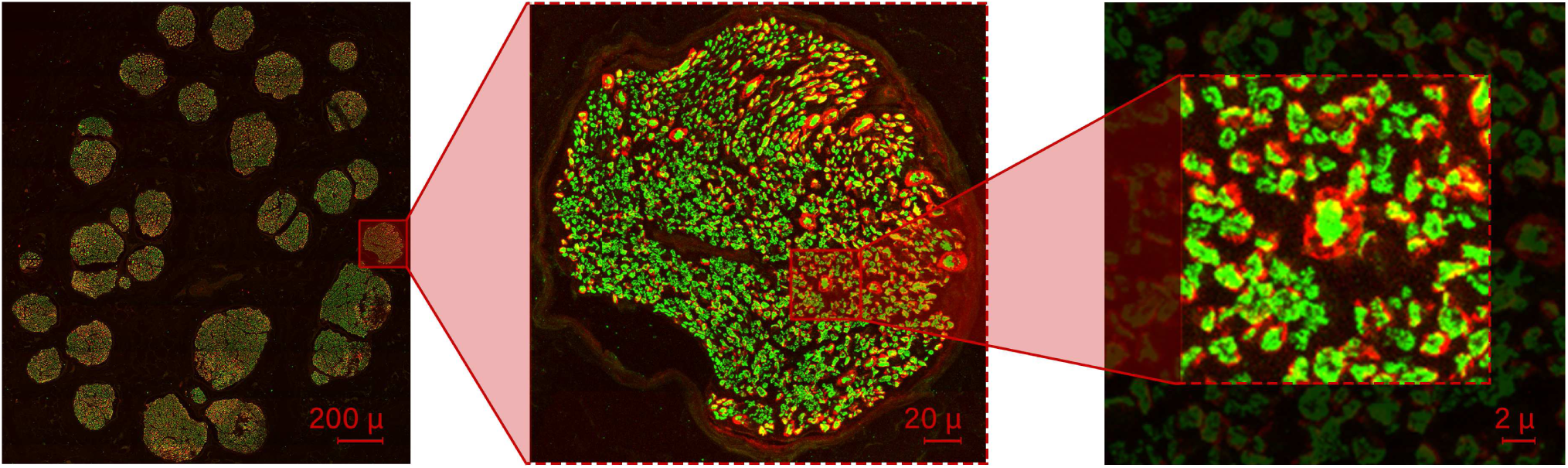
IHC images of the swine vagus nerve used to apply and evaluate the algorithm. Green color indicates the neurofilament (axons) and red color indicates the myelin sheath. The nerve was comprised of 27 fascicles (left panel) and each fascicle was isolated, cropped (center panel) and processed individually. The zoomed image of a smaller region of the nerve (right panel) showcases the variability of the axon sizes, shapes and the presence of myelin in a subset of these axons, along with some cases of overlapping myelin.

First, the algorithm detects individual axons by clustering continuous blobs of green pixels (Fig. 2). Two model parameters govern this operation: a threshold value on the brightness of green pixels from which the clustering can start, and another brightness threshold on the pixels the cluster can spread to. The first threshold was set to 0.5 (out of 1), the second to a lower 0.2, which avoids pale blobs and pale pixel bridges between fibers. Clusters smaller than 10 pixels are discarded as noise.

**Figure. 2.**
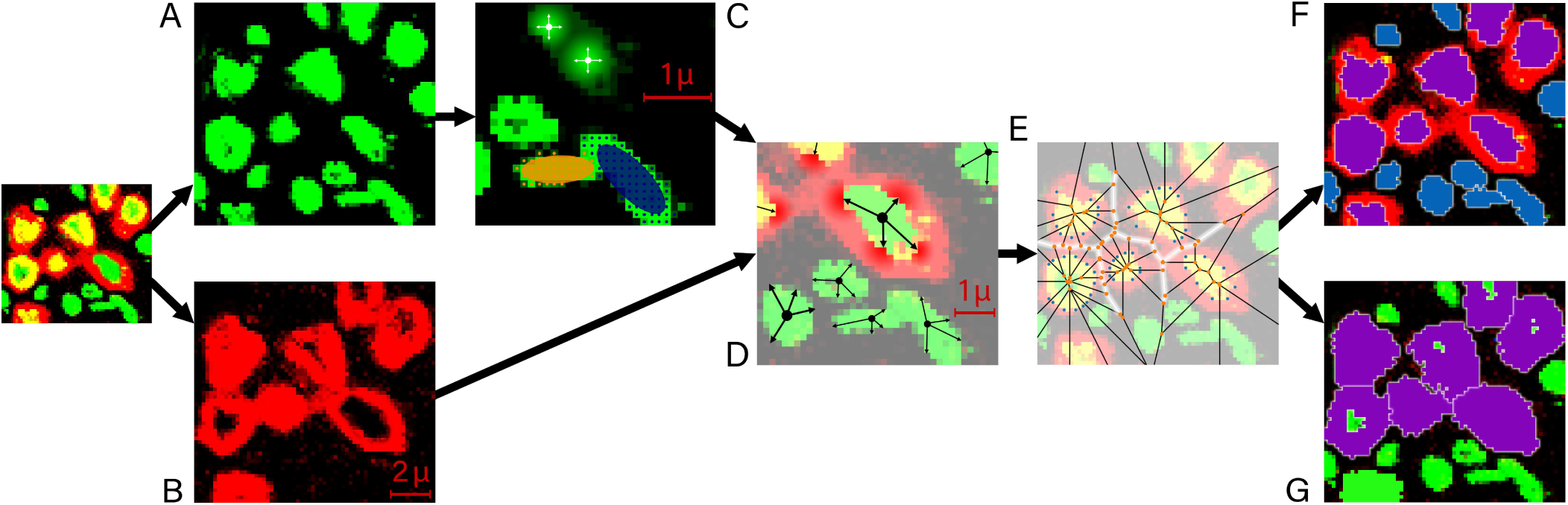
Step-by-step illustration of the segmentation procedure. First, neurofilament (green) and myelin sheath (red) channels are separated (A and B). The algorithm identifies connected neurofilament pixels and clusters them together, forming green homogeneous shapes (top of panel C). Gaussian ellipses are subsequently fitted on the derived shapes to separate potentially overlapping axons (bottom of panel C). Channels are then combined again in order to assign myelin sheaths to corresponding neurofilament shapes by projecting lines from the centroid of the shape outwards until they collide with a black (void) or red (myelin) pixel (D). The more myelin pixels are encountered, the higher probability we assign to the axon being myelinated. Continuous myelin segments are clustered together and assigned to their corresponding axons. However, occasionally, the sheaths of some axons overlap. To distribute overlapping myelin segments between fibers, we performed a Voronoi decomposition on the pixel space using the convex hull of the neurofilament shape as generating points (E). Due to the segmentation process, distinct axons and their myelin sheaths can be differentiated (F and G).

Increasing number of ellipses are fitted to each cluster to separate potentially overlapping axons, which overlap is the residue of the previously mentioned maximum intensity projection. Fitting of a maximum of 5 ellipses is accomplished using Gaussian Mixture Models [12]. The algorithm measures the improvement on the log-likelihood of the data (neurofilament pixel coordinates) when adding one more ellipse to describe the blob. Empirical tests on composite fiber shape separation lead to the improvement thresholds of 0.1, 0.08, 0.06 and 0.04, in correspondence to covering the cluster using 2, 3, 4 and 5 Gaussian ellipses, respectively.

Once individual fibers are identified, the algorithm assigns myelin to fibers when present. Rays from the centroid of the axon are projected outwards in random directions, measuring the color of passed pixels. The algorithm counts the number of times a red pixel or a black pixel is encountered beyond the boundaries of the neurofilament. If at least 20% of the rays terminate in red pixels as opposed to black (void) or green pixels of other axons, the fiber is classified as myelinated and the found myelin pixels are assigned to it. Similar to the process of axon detection, we further cluster continuous sections of myelin starting from the assigned pixels.

In IHC images, myelin clusters of different axons may overlap. To segment touching myelin sheaths, we perform Voronoi decomposition [13] using the green pixels of the neurofilament as generating points – we further optimize the decomposition by only including the pixels along the convex hull of the neurofilament as generating points. Around each point, a Voronoi region is created. Myelin residing within the boundaries of the union of such regions are assigned to the corresponding axon from which the generating pixels are taken.

### C. Feature Extraction

Once segmented, a set of features are derived for each fiber (Fig. 3). These features include the longer and shorter diameters, *d*_*l*_ and *d*_*s*_, the area, *a*, and perimeter of the axon, *p*, in addition to the thickness of the myelin, *t*, and the polar coordinates, radius *r* and angle *α*, of the fiber within the fascicle (Fig. 3).

**Figure 3.**
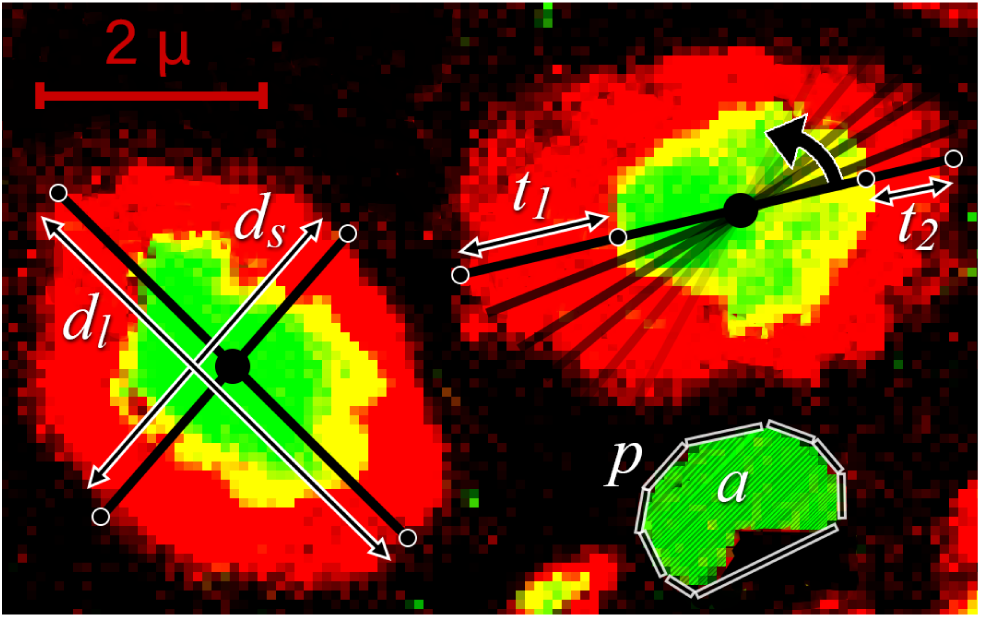
Depiction of some of the single-axon level features that the algorithm extracts following the segmentation and classification procedure. The features include the longer and shorter diameters (*d*_*l*_ and *d*_*s*_), the area (*a*), and perimeter of the axon (*p*), and the thickness of the myelin (*t*_*1*_ and *t*_*2*_ here). The diameter values and the myelin thickness are derived by rotating a line fixated at the centroid of the axon, taking multiple distance measurements at points of intersections with the convex hull of the neurofilament and the myelin sheath. The estimates of the algorithm for this particular example were: *d*_*l*_ = 4.20 µm, *d*_*s*_ = 3.08 µm, *a* = 1.61 µm^2^, *p* = 5.31 µm, *t*_*1*_ = 1.51 µm and *t*_*2*_ = 0.77 µm.

The diameter is computed by forming a polygon out of the convex hull of the fiber (either including the myelin sheath or not), fixating a line at the centroid of the axon, and iteratively rotating it, while registering the distance between the intersecting points of the line and the polygon. The algorithm takes the median, the 5th and 95th percentile of these distance measures to determine the overall, shorter and longer diameter lengths. Myelin thickness is derived in a similar manner: convex hulls of both the neurofilament and the myelin sheath are calculated, before the centroid-fixated line is rotated. At each iteration, two distance measurements are made between the 2-2 corresponding points of neurofilament and myelin polygons that intersect with the line.

## III. Results

We developed a computer vision algorithm that performs fiber-level segmentation of IHC scans of peripheral nerves. The accuracy of the algorithm was tested on images of the swine vagus. Once images were segmented, we extracted features, such as the area, diameter, perimeter of neural fibers, the thickness of myelin sheaths and locational information that can be further processed to generate higher-level structural representations of fascicles. For the example shown in Fig. 3, the values of these features are the following: *d*_*l*_ = 4.20 µm, *d*_*s*_ = 3.08 µm, *a* = 1.61 µm^2^, *p* = 5.31 µm, *t*_*1*_ = 1.51 µm and *t*_*2*_ = 0.77 µm.

We assessed the accuracy of our segmentation model by comparing the segmentation results with the corresponding manually annotated images. Specifically, we assessed the fiber detection and myelination classification performance of the algorithm. An example of manually annotated and corresponding algorithmically derived portion of a fascicle is shown in Fig. 4A and 4B, respectively.

**Figure 4.**
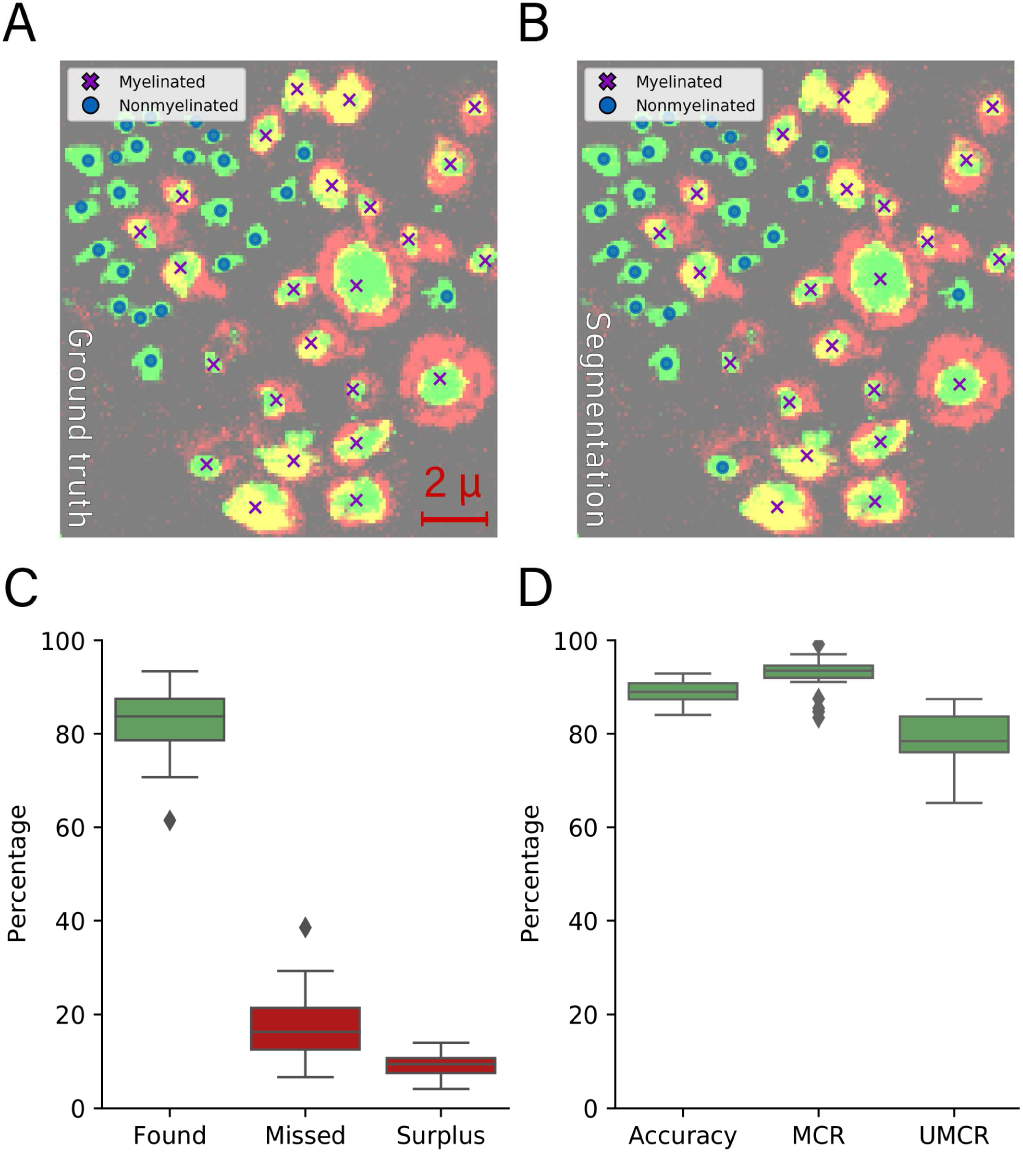
Ground truth manual annotation (A) and corresponding algorithmically derived segmentation (B) of a small nerve section, along with the performance characteristics of the segmentation algorithm. On average our approach identified 82% of the axons, missing 18% and producing a 9% surplus. Once found, the average classification accuracy of fiber myelination averaged at 89%, detecting myelinated fibers (MCR) at a 93%, unmyelinated axon at a 79% accuracy across fascicles

In terms of detection accuracy, we differentiated between the ratio of found, missed and surplus fibers (Fig. 4C). Found fibers were characterized as segmented axons that successfully matched with the annotated ones. The missed fibers are the annotated ones that the algorithm failed to identify. The percentage reported in Fig. 4C is calculated as the number of the found and missed fibers over the total number of annotated fibers. The surplus axons are leftover segmentations of the model, which could not be paired with actual annotated fibers; they are normalized for Fig. 4C with the total number of algorithmically detected fibers. We applied the algorithm on 27 fascicles and calculated the described three accuracy metrics for each fascicle. Our approach successfully discovered an average of 82% of the fibers (with a standard deviation of 7%), missed an average of 18% (SD of 7%) and produced a 9% surplus (SD of 2%) on average across fascicles.

With regards to myelination classification, we evaluated the overall classification accuracy of the model in identifying myelinated versus unmyelinated axons (Fig. 4D). We further computed the myelinated classification ratio (MCR) and unmyelinated classification ratio (UMCR) as the number of successfully identified myelinated axons over the number of annotated myelinated fibers, and the total of correctly classified unmyelinated axons over the number of annotated unmyelinated fibers respectively, to assess the accuracy of successfully identifying myelinated and unmyelinated fibers.

We achieved an overall average accuracy of 89% (SD of 2%), an MCR of 93% (SD of 3%) and a UMCR of 79% (SD of 6%) on average across fascicles.

## IV. Discussion

We developed a novel, automated algorithm to segment and quantify fiber composition in IHC images of a peripheral nerve. The algorithm successfully identified single axons and extracted features such as the diameter of the axon, surface area of neurofilament that each axon expresses, the thickness of myelin surrounding the axon, as well as the relative location of each axon inside a fascicle. The performance of the algorithm was assessed using manually annotated fascicles and was consistently accurate over 80% of the times.

The algorithm can automatically extract highly accurate features in minutes for small, in an hour for large nerves or fascicles. These features provide quantitative descriptions of the anatomy of nerves at an unprecedented, single-axon scale, and is bound to advance a better understanding of the morphological characteristics and variability of peripheral nerve anatomy, across fascicles in the same nerve and across nerves in different subjects.

The features generated by this algorithm can be used to construct computational models of nerves that are biologically accurate. These models could be used to simulate stimulation and recording aspects of nerve interfaces [14]. These simulations can then be used to optimize parameters for selective nerve stimulation and provide insight into possible sources for signals recorded at the surface of nerves using cuff electrodes [15]–[17].

One of the limitations of our current approach is that feature extraction parameters were optimized using just the example of the swine cervical vagus nerve. Even though this nerve was chosen specifically because of its complexity in fiber composition and fascicular anatomy, the algorithm can be made indifferent to the type of nerve or animal species, or to the imaging device. A future version of the algorithm could include implementation of an optimization procedure (e.g. particle swarm), which will further increase accuracy by tuning model parameters to fit the manually annotated images.

